# HOMOCYSTEINE POTENTIATES AMYLOID β-INDUCED CEREBRAL ENDOTHELIAL CELL APOPTOSIS, BLOOD BRAIN BARRIER DYSFUNCTION AND ANGIOGENIC IMPAIRMENT

**DOI:** 10.1101/2023.07.06.547994

**Authors:** Ashley Carey, Rebecca Parodi-Rullan, Rafael Vazquez-Torres, Elisa Canepa, Silvia Fossati

## Abstract

Cerebrovascular dysfunction has been implicated as a major contributor to Alzheimer’s Disease (AD) pathology, with cerebral endothelial cell (cEC) stress promoting ischemia, cerebral-blood flow impairments and blood-brain barrier (BBB) permeability. Recent evidence suggests that cardiovascular (CV)/cerebrovascular risk factors, including hyperhomocysteinemia (Hhcy), exacerbate AD pathology and risk. Yet, the underlying molecular mechanisms for this interaction remain unclear. Our lab has demonstrated that amyloid beta 40 (Aβ40) species, and particularly Aβ40-E22Q (vasculotropic Dutch mutant), promote death receptor 4 and 5 (DR4/DR5)-mediated apoptosis in human cECs, barrier permeability and angiogenic impairment. Previous studies show that Hhcy also induces EC dysfunction, but it remains unknown whether Aβ and homocysteine function through common molecular mechanisms. We tested the hypotheses that Hhcy exacerbates Aβ-induced cEC DR4/5-mediated apoptosis, barrier dysfunction, and angiogenesis defects. This study was the first to demonstrate that Hhcy specifically potentiates Aβ40-E22Q-mediated activation of the DR4/5-mediated extrinsic apoptotic pathway in cECs, including DR4/5 expression, caspase 8/9/3 activation, cytochrome-c release and DNA fragmentation. Additionally, we revealed that Hhcy intensifies the deregulation of the same cEC junction proteins mediated by Aβ, precipitating BBB permeability. Furthermore, Hhcy and Aβ40-E22Q, impairing VEGF-A/VEGFR2 signaling and VEGFR2 endosomal trafficking, additively decrease cEC angiogenic capabilities. Overall, these results show that the presence of the CV risk factor Hhcy exacerbates Aβ-induced cEC apoptosis, barrier dysfunction, and angiogenic impairment. This study reveals specific mechanisms through which amyloidosis and Hhcy jointly operate to produce brain EC dysfunction and death, highlighting new potential molecular targets against vascular pathology in comorbid AD/CAA and Hhcy conditions.

## Introduction

Alzheimer’s Disease (AD), the most common age-associated dementia, is the 6th leading cause of US deaths, currently affecting more than 5.8 million individuals with case numbers exponentially rising (“2020 Alzheimer’s disease facts and figures,” 2020). Pathologically, AD is defined by amyloid beta (Aβ) plaque deposition and neurofibrillary tangles of hyperphosphorylated tau, neuroinflammation, cerebrovascular dysfunction and neuronal death, resulting in cognitive decline (DeTure & Dickson, 2019). Cerebral Amyloid Angiopathy (CAA), a condition involving Aβ deposition (particularly Aβ40 species) around and within the cerebral vasculature, is evident in 85-95% of AD patients and is also common in the aging population (Cortes-Canteli & Iadecola, 2020; DeTure & Dickson, 2019; Fossati et al., 2010). CAA pathology contributes to focal ischemia, cerebral blood flow (CBF) and blood brain barrier (BBB) impairments, as well as cerebral hemorrhages (DeTure & Dickson, 2019). Aβ40 is the main component of CAA deposits, and the Aβ40-E22Q Dutch familial mutant, containing a mutation of glutamic acid to glutamine on amino acid 22 –which accelerates the peptide oligomerization–, is highly associated with early onset CAA, encompassing hemorrhagic strokes and dementia in hereditary cerebral hemorrhage with amyloidosis-Dutch type (HCHWA-D) (Fossati et al., 2010; Levy et al., 1990). In patients with the Dutch mutation, the effects of CAA are accelerated and exacerbated, with disease onset happening already in the 3^rd^ or 4^th^ decade of life. Hence, the Aβ40-E22Q Dutch familial mutant, more aggregation prone and aggressive than Aβ40-WT (Fossati et al., 2010), constitutes a useful tool to study the effect of toxic Aβ aggregates on the cerebral vasculature in experimental conditions.

Recent evidence suggests that cerebrovascular disturbances may precede Aβ-mediated pathology in AD (Korte, Nortley, & Attwell, 2020; Nortley et al., 2019), shifting the focus of the dementia field to understanding the pathological contribution of cardiovascular (CV) factors to AD onset and progression (Iturria-Medina et al., 2016; Love & Miners, 2016). Indeed, it has been demonstrated that CV risk factors can promote AD pathology and increase individuals’ AD risk (Carey & Fossati, 2023; Love & Miners, 2016).

Multiple CV risk factors have been shown to cause endothelial cell (EC) dysfunction leading to cerebral hypoperfusion, which is an early contributor to AD pathology (Di Marco et al., 2015) and BBB abnormalities (Wong et al., 2019). Hyperhomocysteinemia (Hhcy), a disorder involving excess plasma homocysteine (Hcy) levels due to vitamin deficiencies and Hcy metabolism disruption, is prevalent in 5-7% of the US population, and this prevalence increases with age (Son & Lewis, 2023). Hhcy is recognized as both a CV risk factor and a risk factor for AD and vascular dementia (Carey & Fossati, 2023; Kamat, Vacek, Kalani, & Tyagi, 2015; Tinelli, Di Pino, Ficulle, Marcelli, & Feligioni, 2019). It is also known to promote oxidative stress, vascular inflammation, and EC dysfunction (Balint, Jepchumba, Gueant, & Gueant-Rodriguez, 2020; Lai & Kan, 2015; Tinelli et al., 2019). Hhcy and amyloidosis both have detrimental effects on ECs function and health and are both thought to cause cerebrovascular damage. Despite the compelling data in favor of Hcy as a modifiable risk factor, the debate regarding the significance of Hcy mediated health effects, and if these are causal for cerebrovascular dysfunction is still ongoing. Moreover, whether the molecular mechanisms through which endothelial damage is caused are overlapping in CAA and Hhcy remains mostly unknown. Identifying specific pathways activated by both Hhcy and Aβ to promote cerebrovascular pathology and understanding whether there is an additive or synergistic activation of the same molecular mechanisms in ECs will be vital for identifying new therapies for these frequently comorbid conditions, which are increasingly common in aging individuals.

Cerebral ECs are connected by tight junction (TJ) proteins (such as occludin and claudin-5), as well as anchor proteins [such as zona occludens-1 (ZO-1)] and cadherin proteins (VE cadherin) (Yu, Ji, & Shao, 2020). Not only are brain ECs vital for the formation of a physical barrier to protect the cerebral environment, but they contribute to the transport in and out of the brain, by regulating influx and efflux, and to the immunological barrier, by preventing immune cell infiltration and neuroinflammation (Yu et al., 2020). BBB dysfunction is a prominent feature of AD pathology, involving disruption of TJs and loss of barrier integrity. Increased BBB permeability leads to the leakage of neurotoxic blood proteins and immune cells, promoting neuroinflammation and oxidative stress. As our lab and others have demonstrated, AβQ22 and other Aβ peptides directly contribute to BBB permeability (Cuevas et al., 2019; Hartz et al., 2012; R. Parodi-Rullan, Ghiso, Cabrera, Rostagno, & Fossati, 2020).

We have also demonstrated that Aβ oligomers activate caspase- and mitochondria-mediated apoptosis in cerebral ECs through their direct binding with the TNF-related apoptosis-inducing ligand (TRAIL) death receptors (DRs), DR4 and DR5 (Fossati, Ghiso, & Rostagno, 2012c). Ligand binding to DR4/5 results in the activation of caspase 8, to initiate the extrinsic apoptotic pathway (Elmore, 2007). Activated caspase 8 cleaves Bid into activated tBid. tBid increases mitochondrial membrane permeability, inducing release of cytochrome C (CytC), which causes caspase 9 activation (Elmore, 2007). Data from our lab and others have shown that Aβ induces caspase 8 activation and CytC release into the cytoplasm in cerebral ECs (Fossati, Ghiso, & Rostagno, 2012a; Solesio et al., 2018; Xu et al., 2001), eventually resulting in caspase 3 activation and, ultimately, apoptotic cell death (Elmore, 2007), and knocking down DR4/5 resulted in 60-70% protection from Aβ-induced EC apoptosis (Fossati et al., 2012c). As the kinetics for these affects are aggregation-dependent, DRs activation occurs more rapidly in Aβ40-E22Q (AβQ22)-treated ECs, although the same pathways are activated in ECs challenged with Aβ40-WT, once it forms high molecular weight oligomers or protofibrils (Fossati et al., 2012c).

Cerebral angiogenesis, the process of forming new blood vessels, is vital for maintenance of homeostasis within the cerebral vasculature and in the brain parenchyma (Beck & Plate, 2009). Typically, cerebral angiogenesis is activated during hypoxia and hypoperfusion as a mechanism to maintain proper CBF (Beck & Plate, 2009). Previous research has demonstrated that Aβ40-WT and AβQ22 are capable of 90-100% inhibition of EC proliferation, disrupting EC response to angiogenic stimuli (Solito et al., 2009). More recently, our lab has demonstrated that AβQ22 has potent inhibitory effects on EC angiogenesis, compared to other Aβ peptides (R. Parodi-Rullan et al., 2020). Aβ may reach very high concentrations in CAA deposits around the vessels, impairing neo-angiogenesis, which is vitally needed as a repair mechanism, especially when brain vessels are injured by cerebrovascular risk factors, such as Hhcy (R. Parodi-Rullan et al., 2020).

Hence, in this study, we aim to understand whether high levels of Hcy potentiate the effects of Aβ on DR-mediated EC death, BBB dysfunction and angiogenic pathways. The goal of this study is to elucidate whether combined exposure of ECs to Aβ and Hcy promotes exacerbated levels of BBB damage by acting on the same molecular and cellular mechanisms. Overall, we hypothesize that Hhcy potentiates Aβ-mediated TRAIL DR-dependent EC apoptosis, BBB permeability, and angiogenic impairment through additive mechanisms, accelerating the progression cerebrovascular pathology and, ultimately, increasing the risk for AD and dementia.

## Materials and Methods

### Cell Culture

Immortalized human cerebral microvascular endothelial cells (HCMEC/D3) were obtained from Babette Weksler (Cornell University)(Fossati et al., 2010). Cells were grown in EBM-2 (Lonza) and supplemented with growth factors (Hydrocortisone, hFGF-B, VEGF, R3-IGF-1, ascorbic acid, hEGF, and GA-1000) and 5% FBS and maintained at 37°C in a humidified cell culture incubator under a 5% CO_2_ atmosphere. Cells were visualized and imaged using the EVOS M5000 Imaging System (Thermo Fisher Scientific). For the TJ staining, primary human cerebral endothelial cells (HCECs) (Sciencell) were utilized and maintained under the same conditions as the HCMECs.

### Aβ40-E22Q Peptide

For treatments, we utilized the genetic variant of the wild-type (WT) Aβ40 peptide containing the E22Q substitution (Aβ40-E22Q), which is the synthetic homolog of the amyloid subunit present in the vascular deposits in sporadic and familial Dutch-AD cases. The peptide was synthesized by Peptide 2.0. Aβ40-E22Q was dissolved to 1 mM in 1,1,1,3,3,3-hexafluoro-2-propanol (HFIP; Sigma, St. Louis, MI, USA), incubated for 24h to breakdown pre-existing β-sheet structures and lyophilized. Peptides were subsequently dissolved in DMSO to a 10mM concentration, followed by the addition of deionized water to 1mM concentration, and further diluted into culture media (EBM-2 (Lonza) with 1% FBS and no growth factors) to the required concentrations for the different experiments.

### Homocysteine

Hcy was weighed and exposed to 0% O_2_ and 5% CO_2_ conditions in a hypoxic chamber (Coy Laboratory Products) located in a 37°C humidified cell culture incubator for 15 minutes to prevent oxidation before cell treatment. Hcy was prepared fresh for each treatment and dissolved in media to obtain a 100mM concentration and further dissolved to obtain a 10mM concentration. The 10mM Hcy solution was filtered, and additional media was added to obtain the required concentration for the different experiments.

### Mouse Models

Male and female Tg2576 mice, a widely used model of cerebral amyloidosis expressing the Swedish mutation (K670N/M671L), and age-matched wild-type littermates were bred internally. Mice were maintained under controlled conditions (∼22°C, and in an inverted 12-hour light-dark cycle, lights on 10 am to 10 pm) with unrestricted access to food and water. The generation of B6;SJL-Tg(APPSWE)2576Kha mice (Tg2576) on a B6;SJL Mixed Background was as described (Hsiao et al., 1996). Tg2576 mice develop parenchymal Aβ plaques at 11-13 months and vascular Aβ deposition (CAA) at 10-11 months of age (Robbins et al., 2006). Starting at 5 months, WT and Tg2576 mice were supplemented with a Hhcy-inducing diet, consisting of high methionine (Hcy precursor; 20g/kg) and low folate (3.2 mg/kg), vitamin B6 (15mg/kg), and vitamin B12 (55ug/kg) (needed to metabolize Hcy). Mice were sacrificed at 13-14 months of age and tissue lysate from the prefrontal cortex was harvested. All experiments and animal protocols were performed according to protocols approved by the Institutional Animal Care and Use Committee of Temple University School of Medicine and conformed to the National Research Council Guide for the Care and Use of Laboratory Animals published by the US National Institutes of Health (2011, eighth edition).

### Western Blot

Evaluation of DR4 (Invitrogen; 32A242), DR5 (Enzo; ALX-210-743-C200), cFLIP (Cell Signaling; CS563435), BCL2 (abcam; ab196495), Bax (Novus; NBPI-88682), ICAM (Invitrogen; MA5407), ZO1 (Invitrogen; 61-7300)), Claudin-5 (Invitrogen; 35-2500 4C3C2), phosphorylated Claudin-5 (abcam; ab172968), phosphorylated VeCadherin (Invitrogen; 44-11446), VEGFR2 phospho Y1175 (abcam; ab194806), and VEGF-A (Proteintech; 19003-1-AP) was performed using WB analysis after electrophoretic separation on 4-12% Bolt Bis-Tris SDS polyacrylamide gels. Evaluation of Bid/tBid (Cell Signaling; CS#2002) and cleaved caspase 3 (Cell Signaling; C#9664) was performed using WB analysis after electrophoretic separation on Novex WedgeWell 4-20% tris-glycine SDS polyacrylamide gels. Normalization was performed using anti-β actin (Millipore; MAB1501) or anti-GAPDH (Cell signaling; 32233). Proteins were electrotransferred to nitrocellulose membranes (0.45 *μ*m pore size; Amersham, Cytiva LifeSciences) at 110 V for 70 min, using towbin buffer, containing 20% (v/v) methanol. Membranes were blocked with 5% non-fat milk in TBST (or 5% BSA in TBST for the phosphorylated proteins) containing 0.1% Tween 20, and subsequently immunoreacted with the respective primary antibodies for each experiment, as well as a monoclonal antibody against Actin (Millipore; MAB1501) for a loading control, followed by incubation with the appropriate anti-rabbit, anti-mouse, or anti-goat secondary antibodies (1/20,000; LICOR). Membranes were developed with the LICOR Odyssey CLx Immunoblot Imager and blots were analyzed with the LICOR Image Studio software.

### Cell Death ELISA

Apoptotic cell death was assessed as formation of fragmented nucleosomes using the Cell Death Detection ELISA^Plus^ kit (Roche Applied Science) according to the manufacturer’s instructions. Briefly, HCMEC were seeded and, after 24h, treated with 25 µM Aβ40-E22Q, 1mM Hcy, or a combination of both, in EBM-2 media supplemented with 1% FBS. Extranuclear DNA-histone complexes were measured with Cell Death Detection ELISA^Plus^ at 405 nm using the SpectraMax i3x Multi-Mode Microplate Reader (Molecular Devices). Results are expressed as percent change compared with untreated control cells.

### Propidium Iodide Necrosis Staining

Necrosis levels were fluorescently assessed using a Propidium Iodide Nucleic Acid Live Cell Stain (Invitrogen). 10,000 cells/well were seeded in a 96 well plate and, following 24h, were treated with 25uM Aβ40-E22Q, 1mM Hcy, or a combination of both for 24h. A stock of 1mg/ml propidium iodide was dissolved in dH_2_0. Following treatment, the media was removed, and the propidium iodide solution was added to each well as a 1:200 dilution in cell culture media. Cells were incubated with the dye for 1h at room temperature in the dark and were imaged with bright field and an RFP filter using an EVOS M5000 imaging system. 4 randomized images were taken per well. The percentage of propidium iodide positive cells was calculated by counting the total number of cells present in each picture and creating a mask for the fluorescent signal to count the number of cells showing propidium iodide fluorescence [% of Propidium Iodide Positive Cells = (# of cells with Propidium Iodide/total number of cells)*100].

### Caspase-3/7, -8, and -9 Activity Assays

Caspase activation was measured by luminescent assays (Caspase-Glo-3/7, -8 or -9, Promega, Madison, WI, USA), in cells treated for 8h with 50uM Aβ40-E22Q, 1mM Hcy, or a combination of both in EBM-2/1% FBS. Briefly, 10,000 cells/well were seeded in white wall/bottom 96-well plates and were treated following 24h. Caspase-Glo reagent was added to the cell cultures resulting in cell lysis, followed by caspase cleavage of the substrate and generation of a luminescent signal produced by the luciferase reaction. After 1hr incubation the signal, proportional to the amount of caspase activity present, was evaluated in a plate-reading luminometer (SpectraMax i3x Multi-Mode Microplate Reader; Molecular Devices). To inhibit nonspecific background activity, the proteasome inhibitor MG-132 was added to the Caspase-Glo 8 and 9 reagent before the experiment as indicated by the manufacturer. In all cases, results are expressed as percent change compared with untreated control cells.

### Cell Event Fluorescent Caspase 3/7 Assay

Cleaved/Active Caspase 3/7 expression was assessed using the CellEvent^TM^ Caspase-3/7 Green Detection Reagent (ThermoFischer Scientific). 10,000 cells/well were seeded in a 96 well plate and were treated with 25uM Aβ40-E22Q, 1mM Hcy, or a combination of both following 24h. Following treatment, the media was removed, and the fluorescent Caspase 3/7 Green Detection Reagent was added to each well at a concentration of 5uM. Cells were incubated with the dye for 1h at 37°C and were imaged with bright field and a GFP filter using an EVOS M5000 imaging system. 4 randomized images were taken per well. The percentage of cleaved caspase 3/7 positive cells was calculated by counting the total number of cells present in each picture and creating a mask for the fluorescent signal to count the number of cells showing active caspase 3/7 fluorescence [% of Caspase 3/7 Positive Cells = (# of cells with cleaved caspase 3/7**/**total number of cells)*100].

### Cytochrome C Immunocytochemistry

Release of CytC from the mitochondria into the cytoplasm was assessed via immunocytochemistry. HCMECs were seeded in 8 well chamber slides (Millipore) coated with Attachment Factor (Cell systems) and, after 24h, treated with 25uM Aβ40-E22Q, 1mM Hcy, or a combination of both for 6h. Following treatment, cells were stained with MitoTracker Red CMXRos (Invitrogen), a dye that enters the mitochondria only if they present a healthy membrane potential, for 30 minutes at 37°C. Cells were then fixed with 4% PFA (BeanTown Chemical) and permeabilized with 0.2% triton-x100 in PBS for 10 minutes at room temperature. After blocking with 3% BSA in PBS for 1 hour, cells were stained with CytC alexaflour488 (1:500) (BD Biosciences; 560263) in 1% BSA in PBS for 1 hour. Following the staining, cells were mounted with DAPI (blue) mounting media (SouthernBiotech; 0100-20) and images were taken with a fluorescent inverted microscope (Nikon Eclipse Ti2) using the NIS Elements AR Analysis 5.360.02 software and NIS Elements AR 5.360.02 image capturing software.

### Cytochrome C ELISA

Release of cytochrome C from the mitochondria into the cytoplasm was assessed by a human cytochrome C Quantikine ELISA kit (Bio-techne R&D Systems) according to the manufacturer’s recommendations. HCMECs were seeded and following 24h treated with 25uM Aβ40-E22Q, 1mM Hcy, or a combination of both for 6h. Mitochondrial isolation was conducted in order to obtain mitochondrial and cytoplasmic fractions. Data is represented as the ratio of cytoplasmic CytC/mitochondrial CytC (percent change from control).

### ECIS Trans-Endothelial Electrical Resistance

Cerebrovascular endothelial barrier formation was assessed using the ECIS Zθ system (Applied Biophysics). All experimental procedures were performed in 8-well ECIS (8WE10+, Applied Biophysics) 40-electrodes-gold plated arrays pre-treated according to the manufacturer’s instructions. A monodisperse solution of CMECs was seeded and monitored for 48h until the electrical resistance reached a plateau at a frequency of 4000 Hz, indicative of barrier formation. At this point, the cell monolayers were treated with 5 or 10 µM Aβ40-E22Q, 1mM Hcy, or a combination of 5 or 10uM Aβ40-E22Q and 1mM Hcy in EBM-2 media containing 1% FBS and followed for 48 h post-treatment. Barrier permeability was assessed as a decrease in barrier resistance at 4000 Hz compared with untreated cells.

### ZO-1 Immunocytochemistry

ZO-1 expression and localization was assessed via immunocytochemistry. Primary HCECs were seeded in 8 well chamber slides (Millipore) coated with collagen I (0.15 mg/ml) and grown until 100% confluence in complete media. Once confluent, cells were exposed to EBM2 media containing 0.25% FBS and bFGF (1:2000) for 24h to promote TJ formation. Cells were then treated with 10uM Aβ40-E22Q, 1mM Hcy, or a combination of both for 48h. Following treatment, cells were fixed with 4% PFA (BeanTown Chemical) for 10 minutes at room temperature and blocked overnight at 4°C in 20mg/ml BSA. Cells were then incubated with ZO-1 primary antibody (Invitrogen; 61-7300) for 1.5h at room temperature (1:100 in 5mg/ml BSA 0.05% triton-x100). Subsequently, cells were incubated with alexaflour 568 goat anti-rabbit secondary antibody (Life Technologies, A11011) for 1h at room temperature in the dark (1:200 in 5mg/ml BSA 0.05% triton-x100). Cells were then stained with Phalloidin alexaflour488 (Invitrogen; A12379) for 30 minutes at room temperature in the dark (1:100 in 5mg/ml BSA). Following the staining, cells were mounted with DAPI (blue) mounting media (SouthernBiotech; 0100-20) and images were taken with a fluorescent inverted microscope (Nikon Eclipse Ti2) and deconvolved with the 64-bit NIS Elements AR Analysis 5.360.02 analysis software and NIS Elements AR 5.360.02 image capturing software. Area of ZO-1 and Phalloidin fluorescence was analyzed with Halo software (Indica Labs).

### MMP2 Activity Assay

The levels of active MMP2 were measured with a MMP2 activity assay (QuickZyme Biosciences) according to the manufacturer’s recommendations. HCMECs were seeded (350,000 cells/well in 6 well plates) and following 24h treated with 25uM Aβ40-E22Q, 1mM Hcy, or a combination of both for 6h. Following treatment, cells were lysed, and protein extracts were stored at -80°C until the assay was conducted.

### Angiogenesis Inhibition Assay

Inhibition of angiogenesis was assessed using Millipore’s Millicell μ-Angiogenesis Inhibition Assay according to the manufacturer’s recommendations. HCMEC suspensions were seeded in the presence/absence of 1uM Aβ40-E22Q, 500uM Hcy, or a combination of both in a Millicell μ-Angiogenesis Slide containing ECMatrix Gel Solution. Sulforaphane was used as a positive control of angiogenesis inhibition. In all cases, tube formation was monitored after 4h by acquiring pictures with an EVOS M5000 imaging system. Capillary branches meeting length criteria were counted from 4 randomized images for each treatment well. Angiogenesis Progression Score was determined by a scale provided from the manufacture (0= lowest progression level and 5= highest progression level).

### ECIS Wound Healing Assay

EC wound healing capability was assessed using the ECIS Zθ system (Applied Biophysics). All experimental procedures were performed in 8-well ECIS (8W1E, Applied Biophysics) gold plated arrays, presenting a single central electrode, pre-treated according to the manufacturer’s instructions. A monodisperse solution of CMECs was seeded and monitored for 48h until the electrical resistance reached a plateau at a frequency of 4000 Hz, indicative of barrier formation. At this point, the cell monolayers were treated with 10 µM Aβ40-E22Q, 1mM Hcy, or a combination of both in EBM-2 media containing 1% FBS. Approximately 1hr-post treatment, a 20 second wound was inflicted to the ECs (60,000 Hertz; amplitude 5V; wound current= 1400uAmps) and wound healing was followed for 48 h post-injury. Impaired wound healing was defined as decreased barrier resistance following injury compared with untreated cells.

### VEGF-A ELISA

Levels of soluble VEGF-A were assessed by a Human VEGF-A Quantikine ELISA kit (Bio-techne R&D Systems), according to the manufacturer’s recommendations. HCMECs were seeded and following 24h treated with 25uM Aβ40-E22Q, 1mM Hcy, or a combination of both for 24h. Following treatment, media was collected and stored at -80°C until the ELISA was conducted.

### VEGFR2 and EEA1 Immunocytochemistry

VEGFR2 expression and localization was assessed via immunocytochemistry with an EEA1 co-stain. HCMECs were seeded in 8 well chamber slides (Millipore) and, after 24h, treated with 25uM Aβ40-E22Q, 1mM Hcy, or a combination of both for 6h. Following treatment, cells were fixed with 4% PFA (BeanTown Chemical). Cells were permeabilized with 0.2% triton-x100 in PBS for 10 minutes at room temperature. Cells were blocked with 3% BSA in PBS for 1 hour and cells were stained with VEGFR2 (Cell Signaling; C#2479; 1:200) and EEA1 (R&D Systems; AF8047; 5ug/mL) primary antibodies in 1% BSA in PBS overnight at 4°C. Cells were then incubated with secondary antibodies (anti-sheep586 (abcam) and anti-rabbit488 (Invitrogen); 1:200) in 1% BSA/PBS with 0.3% triton-x100 for 2 hours at room temperature. Following the staining, cells were mounted with DAPI (blue) mounting media (SouthernBiotech; 0100-20) and images were taken with a fluorescent inverted microscope (Nikon Eclipse Ti2) and modified with the 64-bit NIS Elements AR Analysis 5.360.02 analysis software and NIS Elements AR 5.360.02 image capturing software. Colocalization of VEGFR2 and EEA1 fluorescence was analyzed using JaCoP (Just another Colocalization) ImageJ plug-in, which creates a mask over the VEGFR2 and EEA1 fluorescence signals and calculates Manders’ coefficients (M1 and M2), which imply the actual overlap of the signals (A over B and B over A, respectively), and represent the true colocalization degree (Zinchuk, Zinchuk, & Okada, 2007). M1 and M2 coefficients were scored from 0 to 1 [e.g., M1=1.0 and M2=0.7, in red (signal A)-green (signal B) pair, indicates that 100% of red pixels colocalize with green, and 70% of green pixels colocalize with red]. The M1 and M2 coefficients values were then multiplied by the percentage area of VEGFR2, accordingly, and plotted.

### Proinflammatory Panel MSD

The levels of proinflammatory cytokine expression was assessed using a V-PLEX proinflammatory panel 1 (human) kit from MesoScale Discovery (MSD) which is able to measure the expression of 10 cytokines per sample. HCMECs were seeded and following 24h treated with 25uM Aβ40-E22Q, 1mM Hcy, or a combination of both for 3h. Following treatment, media was collected and stored at -80°C until the MSD was conducted. The MSD was carried out according to the manufacturer’s recommendations and the protein concentration was measured to normalize each sample.

### Actin Polymerization Assay

The capability of cells to polymerize actin was assessed with an Actin Polymerization/Depolymerization Assay Kit (Abcam) according to the manufacturer’s recommendations. HCMECs were seeded and following 24h treated with 10uM Aβ40-E22Q, 1mM Hcy, or a combination of both for 48h. Following treatment, cells were lysed in a non-denaturing buffer containing 20mM Tris-HCl (pH 7.5) and 20mM NaCl and homogenized with a glass dounce and cell extracts were immediately utilized for the assay. The fluorescence of each sample was read kinetically for 1h and the change in fluorescent signal was calculated by subtracting the final read from the initial read. The percentage activation effect was calculated based on the kits recommendations (percentage activation effect= (ΔSample/ΔPositive Control)*100).

### Statistical Analysis

All experimental graphs are representative of at least 3 independent experiments with 2 or more technical duplicates. Data are represented as means±SEM. Statistical significance was assessed by one-way ANOVA followed by Tukey’s post-hoc test using GraphPad Prism 9. Statistically significant differences required a p value ≤0.05.

## Results

### Hhcy potentiates Aβ-mediated activation of the DR4 and DR5-mediated extrinsic apoptotic pathway in HCMECs

Aβ40 and its vasculotropic variants have previously been shown to activate TRAIL DR-mediated apoptosis in human cerebral microvascular ECs. In particular, we have shown that Aβ40-E22Q upregulates DR4 and DR5 expression and induces activation of the DRs-mediated extrinsic apoptotic pathway, similarly but more rapidly than Aβ40-WT (Fossati et al., 2012c). Hcy has been previously demonstrated to promote cell death in peripheral ECs through DRs, such as Fas (Tyagi, Ovechkin, Lominadze, Moshal, & Tyagi, 2006) (Suhara et al., 2004). However, it’s currently unknown whether Hcy activates TRAIL DR-mediated apoptotic pathways in cerebral endothelial cells. To determine whether high Hcy triggers EC death through activation of the same DR4/5-mediated apoptotic pathway and whether combined challenge of HCMECs with Aβ40-E22Q and Hcy potentiates the activation of this specific apoptotic pathway, we conducted a time course analysis of DR4 and DR5 protein expression, which is known to be associated to their activation (Micheau, Solary, Hammann, & Dimanche-Boitrel, 1999; Poh, Huang, Hirpara, & Pervaiz, 2007; Rossin, Derouet, Abdel-Sater, & Hueber, 2009). HCMECs were treated with 25uM Aβ40-E22Q, 1mM Hcy (concentrations previously shown to induce apoptosis in ECs)(Fossati, Ghiso, & Rostagno, 2012b; Suhara et al., 2004), or a combination of the two for 3h, 8h, and 24h. We observed that Aβ40-E22Q and Hcy upregulated DR4/5 expression at different time points. Combined treatment with Hcy and Aβ40-E22Q potentiated DR4 and DR5 upregulation, resulting in a significant increase compared to control cells at 3h for DR4 **(Fig. 1A)** and at 8h for DR5 **(Fig. 1B).** These results suggest that Hcy promotes early activation of both DR4 and DR5 and that a combined challenge of HCMECs with Aβ40-E22Q and Hcy additively upregulates these DRs expression, with DR4 upregulation occurring earlier in time than DR5.

**Figure 1.**
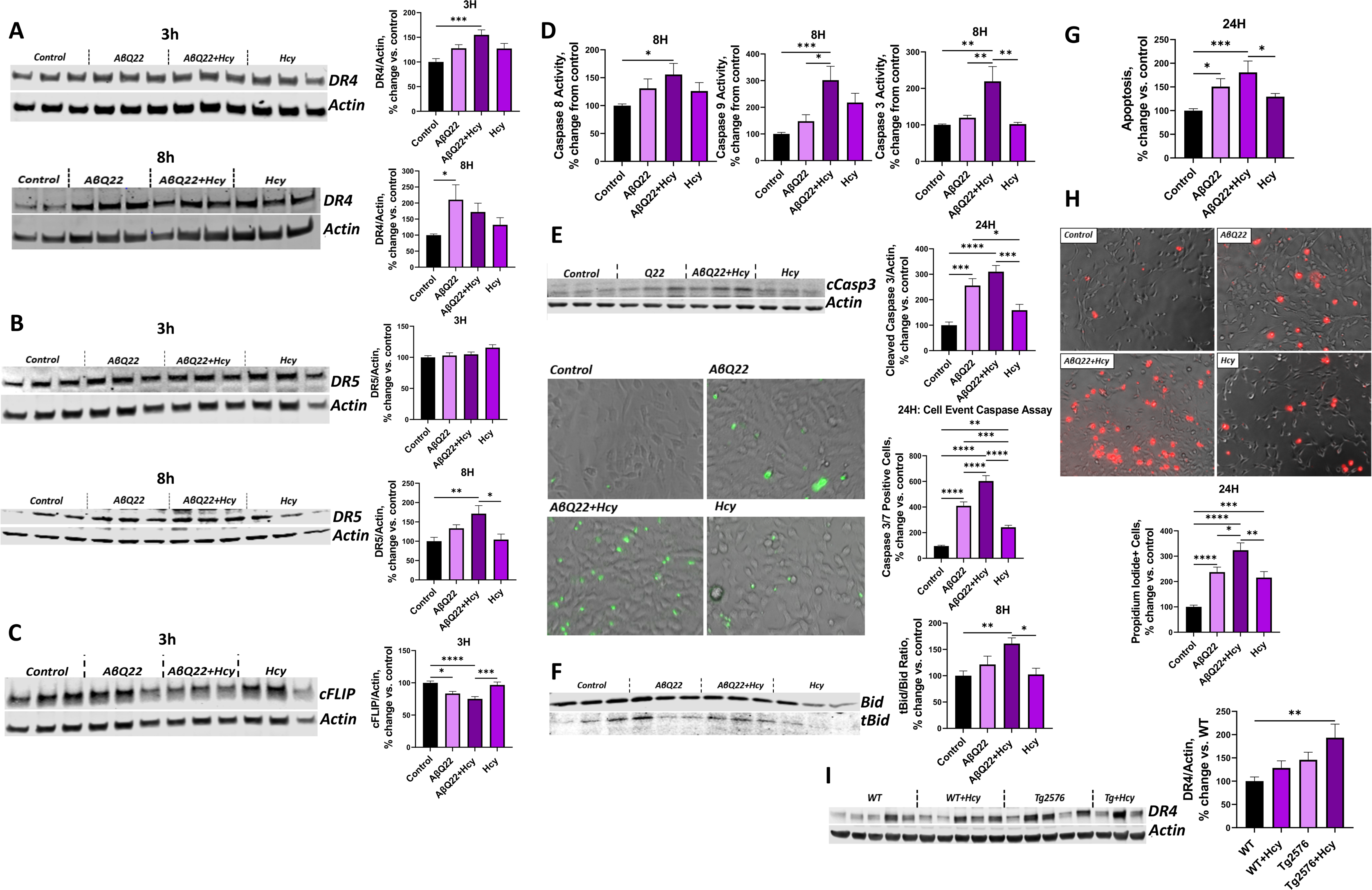
Hcy exacerbates Aβ40-E22Q-mediated DR4 and DR5 upregulation and potentiates activation of the extrinsic apoptotic pathway in HCMECs. (**A/B/C**) HCMECs were treated with 25uM AβQ22, 1mM Hcy, or a combination of the two for 3h or 8h (for DR4 and DR5) and 3h (for cFLIP). (**A**) DR4, (**B**) DR5, and (**C**) cFLIP protein expression was evaluated via western blot. Actin was used for normalization and data is represented as % change vs. control. N ≥ 3 experiments with 2 or more technical replicates; one-way ANOVA and Tukey’s post-test. (**D**) HCMECs were treated with AβQ22, 1mM Hcy, or a combination of the two for 8h. Caspase 8, Caspase 9, and Caspase 3/7 activity was assessed via Promega’s Caspase 3/7, 8, and 9 Glo Assays. Data is represented as % change vs. control. (**E**) HCMECs were treated with 25uM AβQ22, 1mM Hcy, or a combination of the two for 24h. Cleaved caspase 3 protein expression was evaluated via western blot. Actin was used for normalization and data is represented as % change vs. control. The amount of cleaved caspase 3/7 present in cells was evaluated with Thermo Fischer’s Cell Event Caspase 3/7 fluorescence assay (imaged with Invitrogen’s EVOS M5000 microscope). The % of caspase 3/7 positive cells was calculated as (caspase 3/7 positive cells/total cell number)*100. Data is represented as % change vs. control. (**F**) HCMECs were treated with 25uM AβQ22, 1mM Hcy, or a combination of the two for 8h. tBid/Bid protein expression ratio was evaluated via western blot and actin was used for normalization. Data is represented as % change vs. control. (**G**) HCMECs were treated with 25uM AβQ22, 1mM Hcy, or a combination of Hcy and AβQ22 for 24h. Apoptosis was assessed as fragmented nucleosomes via a Cell Death ELISA (CDE). Data is represented as % change vs. control. (**H**) HCMECs were treated with 25uM AβQ22, 1mM Hcy, or a combination of the two for 24h. The amount of propidium iodide positive cells was evaluated with a live cell propidium iodide stain imaged with Invitrogen’s EVOS M5000 microscope and the % of propidium iodide positive cells was calculated as (propidium iodide positive cells/total cell number)*100. Data is represented as % change vs. control. For cell experiments in A-H, N= 3 experiments with 2 or more technical replicates; one-way ANOVA, Tukey’s post-test. (**I**) Western blot analysis of DR4 protein Expression in WT and Tg2576 mice -/+ Hhcy. Actin was used for normalization and data is represented as % change vs. control. N= 3 or more mice per group; n=2 technical replicates. One-way ANOVA, Tukey’s post-test. (**** p<0.0001, *** p<0.001, ** p<0.01, * p<0.05).

Activation of DRs, such as DR4 and DR5, directly induces caspase 8 activation (Cullen & Martin, 2009). cFLIP is an endogenous protein inhibitor of caspase 8 (Kataoka, 2005). Thus, a downregulation in cFLIP promotes caspase 8 activation, allowing for the progression of the extrinsic apoptotic pathway. To determine whether Hcy itself downregulates cFLIP expression and whether combined challenge with Aβ40-E22Q and Hcy additively decreases cFLIP, HCMECs were challenged with 25uM Aβ40-E22Q, 1mM Hcy, or a combination of the two for 3h. cFLIP protein expression was evaluated via western blot. Hcy itself did not significantly downregulate cFLIP expression. However, Hcy co-treatment potentiated the downregulation of cFLIP-induced by Aβ40-E22Q in HCMECs **(Fig. 1C)**.

Active caspase 8 promotes the cleavage of Bid into activated tBid, which is responsible of the recruitment of the intrinsic, mitochondria-mediated apoptotic pathway, resulting in mitochondrial CytC release, caspase 9, and eventually caspase 3 activation (Fossati et al., 2012c). To determine whether Hcy promotes caspase 8, 9 and 3 activation within cerebral ECs and whether combined challenge of HCMECs with Aβ40-E22Q and Hcy potentiates activation of these caspases, we conducted luminescent caspase-glo activity assays. HCMECs were treated with Aβ40-E22Q, Hcy, or a combination of the two for 8h. HCMECs treated with Aβ40-E22Q and Hcy showed a significant caspase 8, 9, and 3 activation, while activation by each challenge alone did not reach significance. Particularly, caspase 3 activation at 8h was synergistically increased by the 2 challenges (**Fig. 1D**). The activation of caspase 3 was still potentiated in HCMECs treated with Aβ40-E22Q and Hcy for 24h, as assessed via both western blot analysis and a fluorescent cleaved caspase 3 assay (**Fig. 1E**). Since activation of caspase 8 induces Bid cleavage into tBid, Bid and tBid protein expression was evaluated by WB in HCMECs treated with 25uM Aβ40-E22Q, 1mM Hcy, or a combination of the two for 8h. The ratio of tBid to Bid was significantly higher in HCMECs treated with both Aβ40-E22Q and Hcy, suggesting that the combination treatment promotes Bid cleavage into tBid (**Fig. 1F**). DNA fragmentation, indicative of the terminal stages of apoptosis, was also measured by the Cell Death ELISA^PLUS^ assay in HCMECs treated with 25uM Aβ40-E22Q, 1mM Hcy, or a combination of the two for 24h. Combined treatment with Aβ40-E22Q and Hcy additively potentiated DNA fragmentation compared to each challenge alone (**Fig. 1G**). Additionally, necrosis was evaluated using a propidium iodide (PI) staining after the same 24h challenges. Treatment of HCMECs with Aβ40-E22Q or Hcy independently increased PI levels compared to control cells and the combined treatment of HCMECs with Aβ40-E22Q and Hcy revealed a significant potentiation of necrosis compared to each challenge alone (**Fig. 1H**).

To confirm *in vivo* whether Hhcy potentiates Aβ-mediated upregulation of DR4, western blot analysis of DR4 expression was performed in the brain of WT and Tg2576 mice previously subjected from 5 to 13 months of age to a Hhcy-inducing diet, or a control diet. WT mice with Hhcy and Tg2576 mice demonstrated a trend towards increased DR4 expression within the pre-frontal cortex, while Tg2576 mice with Hhcy showed a significant upregulation of DR4, confirming the additive effects observed *in vitro* (**Fig. 1I**).

### Hhcy exacerbates Aβ-mediated mitochondrial apoptotic mechanisms in HCMECs

Aβ40-E22Q was previously shown to promote release of CytC from the mitochondria to the cytoplasm in cerebral ECs (Fossati et al., 2010; Solesio et al., 2018). Hcy has also been found to promote CytC release from peripheral ECs (Tyagi et al., 2006). To determine whether Hcy alone promotes CytC release in cerebral ECs and to assess whether combined challenge with Aβ40-E22Q and Hcy potentiates CytC release, we conducted CytC immunocytochemistry and ELISA assays after 6h treatments. Following treatment, HCMECs were stained with DAPI (blue), MitoTracker Red CMXRos (red), and CytC (green). CytC staining in control HCMECs highlighted, as expected, chain-like mitochondrial structures, which colocalized with MitoTracker Red CMXRos, a dye that enters the mitochondria upon a healthy membrane potential **(Fig. 2A)**. HCMECs treated with Aβ40-E22Q demonstrated a partial loss of CytC chain structure, perinuclear localization of mitochondria, and a loss of mitochondrial membrane potential in some of the cells. Similar effects were induced by treatment with Hcy **(Fig. 2A)**. However, when HCMECs were treated with a combination of Aβ40-E22Q and Hcy, a complete loss of mitochondrial CytC chain structure was observed, with significant evidence of mitochondrial fragmentation and most of the CytC signal was lost, suggesting that CytC had been more rapidly released from the mitochondria into the cytoplasm **(Fig. 2A)**. To confirm this phenomenon, following treatment, a mitochondrial isolation was conducted to obtain mitochondrial and cytoplasmic fractions from HCMECs treated with Aβ40-E22Q, Hcy, or the combination, for 6h. For each sample, CytC levels were measured in mitochondrial and cytoplasmic fractions using a CytC ELISA and the ratio of CytC in the cytoplasm vs CytC in the mitochondria was calculated, with a higher ratio being indicative of more cytoplasmic CytC and less mitochondrial CytC. HCMECs treated with either Aβ40-E22Q or Hcy alone demonstrated trends towards increased CytC release into the cytoplasm, but when ECs were treated with the combination of Aβ40-E22Q and Hcy, a significantly higher CytC release was observed **(Fig. 2B)**. Bax and BCL2 protein expression, whose equilibrium mediates mitochondrial voltage-dependent anion channel opening and CytC release, were measured in the same conditions. HCMECs treated with Aβ40-E22Q for 6h demonstrated a significant upregulation of the pro-apoptotic protein Bax, while HCMECs treated with Hcy demonstrated a trend towards increased Bax protein expression **(Fig. 2C)**. In HCMECs treated with a combination of Aβ40-E22Q and Hcy, however, the increase in Bax expression was the most significant compared to control conditions **(Fig. 2C)**. Additionally, BCL2 expression was significantly decreased in HCMECs treated with a combination of Aβ40-E22Q and Hcy for 6h **(Fig. 2C)**.

**Figure 2.**
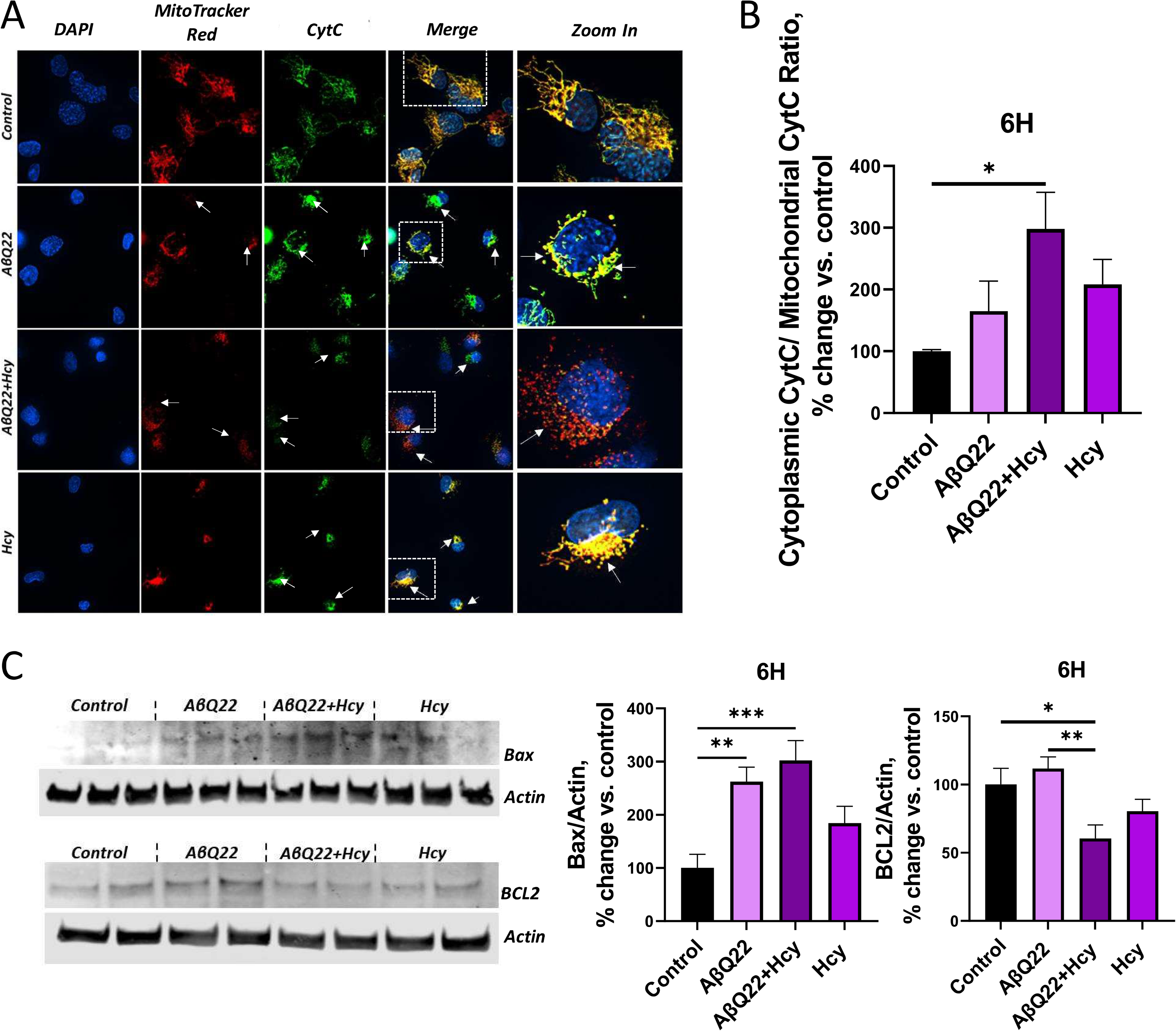
Aβ40-E22Q+Hcy treatment potentiates cytochrome C release, pro-apoptotic Bax overexpression, and decreased anti-apoptotic BCL2 expression in HCMECs. (**A/B**) HCMECs were treated with 25uM AβQ22, 1mM Hcy, or a combination of the two for 6h and Cytochrome C (CytC) release was evaluated by immunocytochemistry (**A**) and ELISA (**B**). (**A**) HCMECs were stained with DAPI (blue), MitotrackerRed (red), and CytC (green). (**B**) Following treatment, mitochondrial and cytoplasmic extracts were obtained and utilized for a CytC ELISA. Data is represented as the ratio of cytoplasmic CytC signal to mitochondrial CytC signal (% change vs. control). (**C**) HCMECs were treated with 25uM AβQ22, 1mM Hcy, or a combination of the two for 6h. Bax and BCL2 protein expression were evaluated via western blot and actin was used for normalization. Data is represented as % change vs. control. For **A-C**, N ≥ 3 experiments with 2 or more technical replicates; one-way ANOVA and Tukey’s post-test. (**** p<0.0001, *** p<0.001, ** p<0.01, * p<0.05).

### Hhcy intensifies Aβ-mediated deregulation of BBB junction proteins and loss of barrier resistance

We have recently shown that Aβ40-E22Q decreases trans-endothelial electrical resistance (TEER) of cerebral EC monolayers (R. Parodi-Rullan et al., 2020). Hcy has also been demonstrated to affect BBB integrity (Beard, Reynolds, & Bearden, 2011; Kamath et al., 2006; Rhodehouse, Mayo, Beard, Chen, & Bearden, 2013). To determine whether combined treatment of HCMECs with Aβ40-E22Q and Hcy additively diminishes barrier resistance, ECIS technology was utilized to monitor TEER over time. Following barrier formation, HCMECs were treated with sublethal concentrations of Aβ40-E22Q (5uM or 10uM), 1mM Hcy, or a combination of the two and TEER was monitored for 48h. Cells treated with Aβ40-E22Q or Hcy separately demonstrated a significant decrease in TEER **(Fig. 3A)**. Moreover, when cells were treated with the combination of Aβ40-E22Q (both the 5 and 10uM) and Hcy, an additive decrease in TEER was observed **(Fig. 3A)**, indicative of an additively increased BBB permeability. To understand whether early junction protein changes are responsible for Hcy and Aβ40-E22Q-induced loss of barrier resistance, we investigated the expression of TJ proteins at early timepoints after challenge. Western blot was utilized to evaluate known mediators of endothelial barrier permeability (Hatanaka, Simons, & Murakami, 2011; Sidibe & Imhof, 2014; Yamamoto et al., 2008), such as phosphorylated VE-cadherin, claudin5, and phosphorylated claudin5 expression, in HCMECs treated with 25uM Aβ40-E22Q, 1mM Hcy, or the combination, for either 3h or 6h. At 3h, the combination of Aβ40-E22Q and Hcy significantly potentiated the increase in phosphorylated VE-cadherin, an adherent protein that upon phosphorylation causes ECs to retract from one another (Hatanaka et al., 2011; Sidibe & Imhof, 2014) **(Fig. 3B)**. Also, after 3h treatment with Aβ40-E22Q and Hcy, ECs revealed a significant reduction in claudin5 expression compared to controls and Aβ treatment alone **(Fig. 3C)**, which was maintained at 6h **(Fig. 3D)**. The combination treatment also induced an additive increase in phosphorylated claudin5 at 6h, which has been associated with the loss of BBB integrity (Yamamoto et al., 2008) **(Fig. 3D)**. To evaluate possible mediators if immune cells adhesion, the expression of ICAM (Intercellular Adhesion Molecule 1), a protein expressed by ECs to promote immune cell extravasation, was also investigated in HCMECs treated with 25uM Aβ40-E22Q, 1mM Hcy, or the combination, for 6h. Cells treated with the combination treatment demonstrated a significant increase in ICAM expression compared to controls **(Fig. 3E)**. Since ICAM upregulation is typically mediated by immune activation through cytokine production, we asked whether cytokine expression preceded ICAM overexpression in our cells. A multiplex cytokine assay (MSD) performed on media from HCMECs treated with Aβ40-E22Q, Hcy, or the combination for 3h, revealed that, while Hcy alone decreased IL2 levels, Aβ40-E22Q alone and the combination of Aβ40-E22Q and Hcy induced a significant increase in IL2 expression, with the combination treatment inducing the most significant increase **(Fig. 3F).** Other pro-inflammatory cytokines tested were not significantly increased at this time point (**Supplementary Fig 1)**. In addition, the expression of MMP2, an enzyme responsible for breaking down extracellular matrix proteins like collagen, was explored in HCMECs treated with Aβ40-E22Q, Hcy, or the combination of the two for 6h (H. Wang et al., 2020). Cells treated with both Aβ40-E22Q and Hcy demonstrated a significant and additive increase in MMP2 expression compared to controls **(Fig. 3G)**. Moreover, we observed a synergistical increase in MMP2 activity when cells were challenged with the combined treatment (**Fig 3H**).

**Figure 3.**
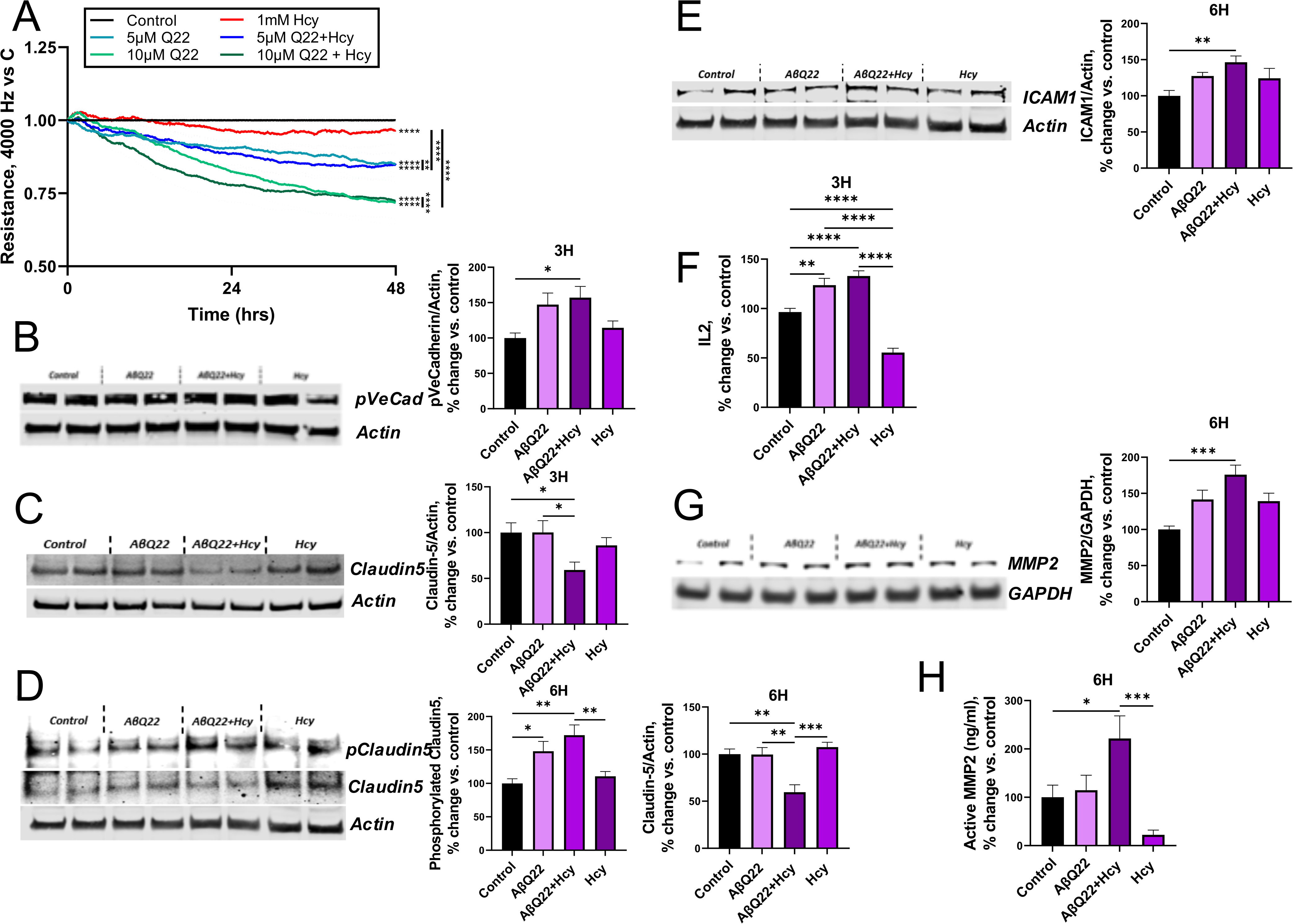
Aβ40E22Q+Hcy challenge potentiates decrease in barrier resistance, tight junction protein dysfunction, and increases in ICAM, MMP2, and IL2. (**A**) Trans-endothelial electrical resistance was monitored for 48h post-barrier formation with the ECIS Zθ system (Applied Biophysics). After formation of a HCMEC-barrier, cells were treated with 5 or 10uM AβQ22, 1mM Hcy, or a combination of the two. Data are represented as resistance change vs. control (black line) (N= 3 experiments with 2 technical replicates per group; one-way ANOVA). (**B**) HCMECs were treated with 25uM AβQ22, 1mM Hcy, or a combination of the two for 3h. Phosphorylated VE-cadherin protein expression was evaluated via western blot and actin was used for normalization. Data is represented as % change vs. control (N ≥ 3 experiments with 2 or more technical replicates; one-way ANOVA; Tukey’s post-test). (**C**) HCMECs were treated with 25uM AβQ22, 1mM Hcy, or a combination of the two for 3h. Claudin-5 protein expression was evaluated via western blot and actin was used for normalization. Data is represented as % change vs. control (N ≥ 3 experiments with 2 or more technical replicates; one-way ANOVA; Tukey’s post-test). (**D**) HCMECs were treated with 25uM AβQ22, 1mM Hcy, or a combination of the two for 6h. Claudin-5 and phosphorylated claudin-5 protein expression was evaluated via western blot and actin was used for normalization. Data is represented as % change vs. control (N ≥ 3 experiments with 2 or more technical replicates; one-way ANOVA; Tukey’s post-test). (**E**) HCMECs were treated with 25uM AβQ22, 1mM Hcy, or a combination of the two for 6h. ICAM protein expression was evaluated via western blot and actin was used for normalization. Data is represented as % change vs. control (N ≥ 3 experiments with 2 or more technical replicates; one-way ANOVA; Tukey’s post-test). (**F**) HCMECs were treated with 25uM AβQ22, 1mM Hcy, or a combination of the two for 3h and media was collected to run a multiplex inflammatory cytokine assay (MesoScale Discovery) and protein concentration was used for normalization. Data is represented as % change vs. control (N ≥ 3 experiments with 2 or more technical replicates; one-way ANOVA; Tukey’s post-test). (**G/H**) HCMECs were treated with 25uM AβQ22, 1mM Hcy, or a combination of the two for 6h. (**G**) MMP2 protein expression was evaluated via western blot and GAPDH was used for normalization. (**H**) MMP2 activity was evaluated with an MMP2 activity assay (QuickZyme Biosciences). Data is represented as % change vs. control (N ≥ 3 experiments with 2 or more technical replicates; one-way ANOVA; Tukey’s post-test). (**** p<0.0001, *** p<0.001, ** p<0.01, * p<0.05).

Overall, these changes in specific functional components of the cerebral endothelial barrier explain the severe loss in TEER observed in HCMECs treated with the combination treatment.

Since a dramatic decrease in TEER was observed at 48h, especially in HCMECs treated with the combination of Aβ40-E22Q and Hcy, the expression of TJ proteins was also assessed in HCMECs treated with 10uM Aβ40-E22Q, 1mM Hcy, or a combination of the two for 48h, to assess the state of the barrier after longer exposure. The deregulation of ZO1 expression and membrane localization was assessed by immunocytochemistry. The cells were treated either prior to endothelial barrier formation (Fig. 4A) or after establishment of the barrier (Fig. 4B). During the process of EC barrier formation, control cells demonstrate a robust network of actin filaments (green) and clear evidence of ZO1 (red) expression between cells attaching to these actin filaments to anchor the cells closer together **(Fig. 4A)**. HCMECs treated with Aβ40-E22Q or Hcy individually demonstrated a derangement in the actin cytoskeleton and appeared to have a decreased connection of ZO1 to the actin filaments **(Fig. 4A)**. HCMECs treated with the combination of Aβ40-E22Q and Hcy demonstrated a loss of ZO1 expression and an even more extreme loss of structure in the actin cytoskeleton **(Fig. 4A)**. Once the barrier was established, control cells showed, as expected, a continuous outline of ZO1 bordering neighbor cells **(Fig. 4B)**. In HCMECs treated with Aβ40-E22Q or Hcy post-barrier formation, noticeable interruptions in the ZO1 outline can be observed, as well as a loss in actin filaments, while HCMECs treated with the combination of Aβ40-E22Q and Hcy revealed an almost complete loss of the ZO1 outline, and a severe disruption of the actin filament network, further corroborated by the quantification of ZO-1 and Phalloidin fluorescence intensity **(Fig. 4C)**. The proper cobblestone EC morphology was also severely altered by the combination treatment. Western blot analysis confirmed that ZO1 expression was significantly reduced by the combination treatment at this time point **(Fig. 4D)**.

**Figure 4.**
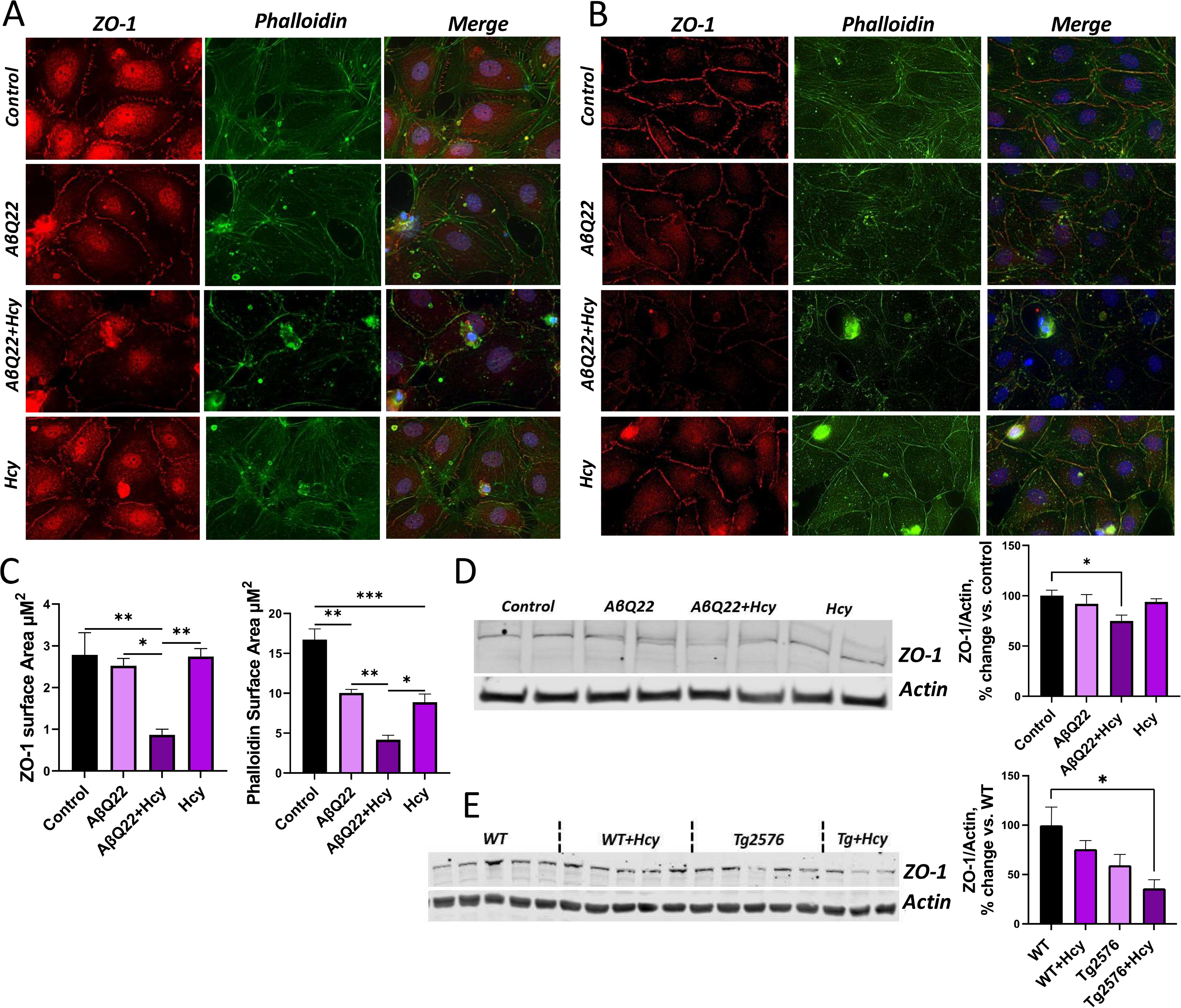
Hhcy potentiates Aβ40E22Q-mediated loss of ZO-1 in human cerebral endothelial cells and in the mouse cortex. (**A/B**) Primary human cerebral endothleial cells were treated with 10uM AβQ22, 1mM Hcy, or a combination of the two either prior to barrier formation (A) or post-barrier formation (**B**). ZO-1 and phalloidin expression were evaluated by immunocytochemistry. Primary human cerebral endothelial cells were stained with DAPI (blue), ZO-1 (red), and phalloidin (green). (**C**) Fluorescence of ZO-1 and phalloidin in panel (B) were quantified with the Halo software (Indica Labs). (**D**) HCMECs were treated with 10uM AβQ22, 1mM Hcy, or a combination of the two for 48h. ZO-1 protein expression was evaluated via western blot and actin was used for normalization. Data is represented as % change vs. control (N ≥ 3 experiments with 2 technical replicates; one-way ANOVA; Tukey’s post-test). (**E**) Western blot analysis was utilized to assess ZO-1 protein expression in WT and Tg2576 mice -/+ Hhcy. Actin was used for normalization and data is represented as % change vs. control (N= 3 or more mice per group; n=1 technical replicate; one-way ANOVA, Tukey’s post-test). (**** p<0.0001, *** p<0.001, ** p<0.01, * p<0.05).

To elucidate *in vivo* whether Hhcy contributes to the loss of ZO-1 and potentiates Aβ-mediated ZO-1 decrease, western blot analysis of ZO-1 expression was evaluated within the pre-frontal cortex of WT and Tg2576 mice with or without Hhcy-inducing diet. WT mice with Hhcy and Tg2576 mice demonstrated a trend towards decreased ZO-1 expression. A significant and additive loss of ZO-1 protein expression was observed in Tg2576 mice with Hhcy (**Fig. 4E**), confirming our *in vitro* results.

### Aβ and Hcy additively decrease angiogenic capabilities of HCMECs

We have recently shown that Aβ peptides, including Aβ40-E22Q, promote deficits in human cerebral EC angiogenesis (R. Parodi-Rullan et al., 2020). In particular Aβ40-E22Q significantly inhibited angiogenesis already at the concentration of 1uM. Hcy has been observed to disrupt angiogenic capabilities of HUVEC ECs; however, it is not established if it can affect cerebral angiogenesis (Pan, Yu, Huang, Zheng, & Xu, 2017; Tian et al., 2020; Zhang et al., 2012). To determine whether Hcy promotes angiogenic deficits in cerebral ECs and whether combined challenge of HCMECs with Aβ40-E22Q and Hcy additively decreases HCMECs angiogenic capability, we performed an angiogenesis inhibition assay. ECs were treated with 1uM Aβ40-E22Q, known as the lowest dose capable of inducing angiogenesis inhibition in this vessel formation assay in our previous studies, 500uM Hcy, or the combination of the two for 4h. Angiogenesis progression score and branch number of cerebral ECs were analyzed. Treatment with Hcy decreased angiogenesis progression score and branch number. When HCMECs were treated with both Aβ40-E22Q and Hcy, an additive decrease in angiogenesis progression score and vessel branch number was observed **(Fig. 5A)**. Since particular cytokines are known to trigger angiogenesis, angiogenesis-related cytokine release was assessed utilizing a multiplex cytokine array from MesoScale Discovery (MSD) as above. HCMECs were treated with Aβ40-E22Q, Hcy, or a combination of the two for 3h and media was collected. The expression of IL13, IL4, IL8, IL1β, and IL6, which all have been previously found to play a role in the promotion of angiogenesis, were additively decreased in HCMECs treated with the combination of Aβ40-E22Q and Hcy **(Fig. 5B)**. Changes in other cytokines not specifically related to angiogenesis are presented in **Supplementary Fig 1.** Moreover, to investigate whether the combination of Aβ and Hcy affects EC repair mechanisms, a wound healing assay was conducted using the ECIS technology. Following barrier formation, HCMECs were treated with 10uM Aβ40-E22Q, 1mM Hcy, or a combination of the two and 1h post-treatment, a circular wound was inflicted in the center of the EC barrier by a strong electrical current. Recovery from the wound, indicated by increases in TEER post-wound, was monitored for 48h. HCMECs treated with Hcy were able to recover up to the control level, while HCMECs treated with Aβ40-E22Q showed impairment in recovery ability, only reaching about half the recovery of the controls **(Fig. 5C)**. HCMECs treated with the combination of Aβ40-E22Q and Hcy were completely unable to recover from the wound, showing significant impairment in wound healing compared to controls and both treatments alone **(Fig. 5C)**. Since the polymerization of actin is essential for cell proliferation, wound healing and endothelial barrier formation and Aβ40-E22Q and Hcy affect the actin cytoskeleton in HCMECs **(Fig. 4B/C)**, the ability of HCMECs to polymerize actin was assessed in cells treated with Aβ40-E22Q, Hcy, or the combination of the two for 48h. HCMECs treated with the Aβ peptide demonstrated a significant decrease in actin polymerization compared to control cells and HCMECs treated with Hcy demonstrated a similar, but non-significant trend **(Fig. 5D)**. As expected, treatment with the combination of Aβ40-E22Q and Hcy resulted in the most significant decrease in actin polymerization capability **(Fig. 5D)**.

**Figure 5.**
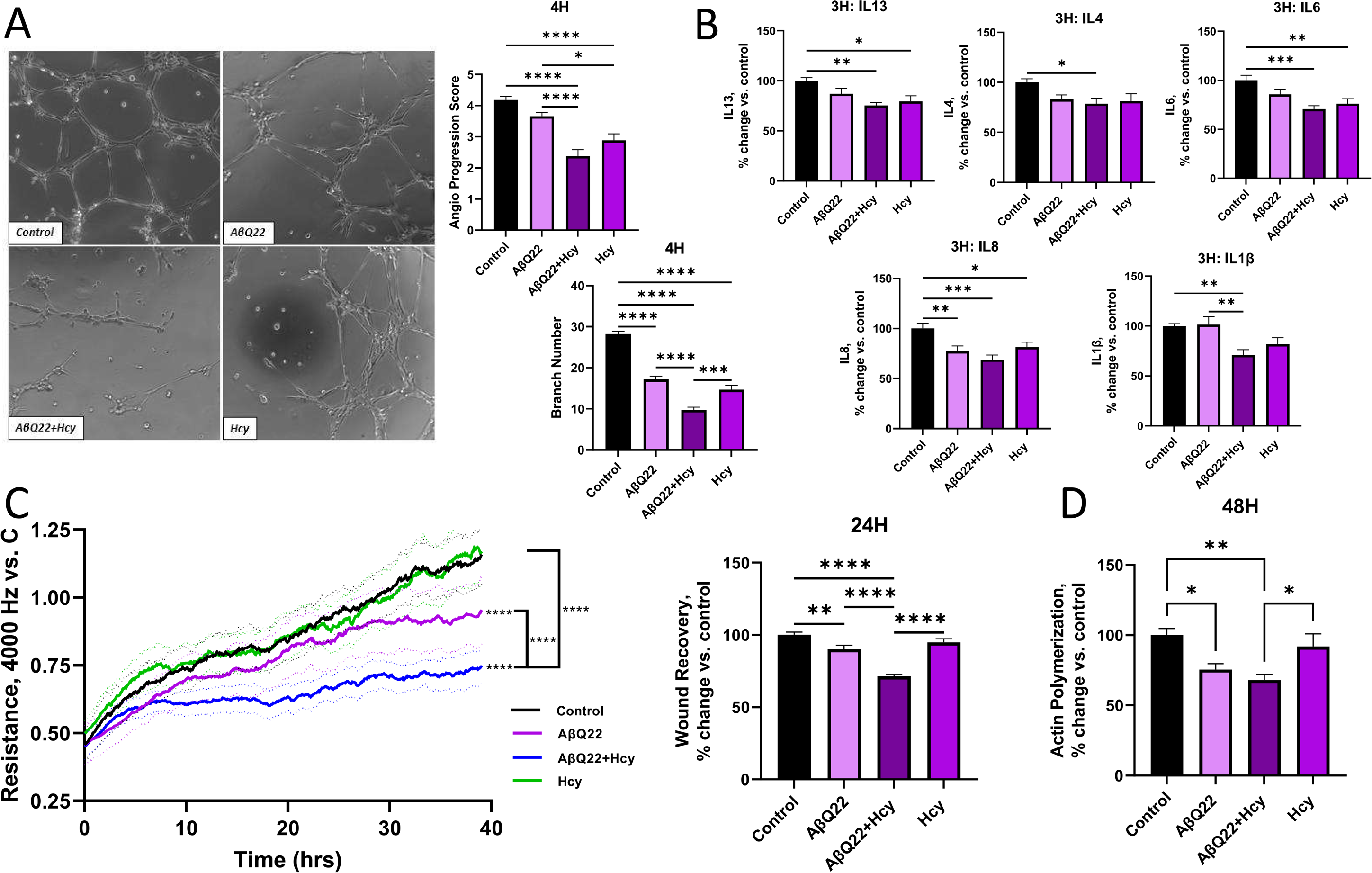
Aβ40E22Q+Hcy potentiates decreases in angiogenic and wound healing capabilities, angiogenesis-related cytokine expression, and actin polymerization in HCMECs. (**A**) HCMECs were treated with 1uM AβQ22, 500uM Hcy, or a combination of the two for 4h. Vessel branch number and angiogenesis progression score were assessed via an angiogenesis inhibition kit. Data is represented as % change vs. control (N= 3 experiments with 2 or more technical replicates; one-way ANOVA, Tukey’s post-test). (**B**) HCMECs were treated with AβQ22, Hcy, or a combination of the two for 3h and media was collected to run a multiplex inflammatory cytokine assay (MesoScale Discovery) and protein concentration was used for normalization. Data is represented as % change vs. control (N ≥ 3 experiments with 2 or more technical replicates; one-way ANOVA; Tukey’s post-test). (**C**) HCMEC would healing ability was assessed with an ECIS would healing assay. Post-barrier formation, HCMECs were treated with 10uM AβQ22, 1mM Hcy, or a combination of the two and 1.5h post-treatment were wounded for 20 seconds (60000hz). Following the wound, HCMEC trans-endothelial electrical resistance was monitored and an increase in TEER post-wound is indicative of wound healing. Data is represented as resistance change vs. control (black line) (N= 4 experiments with 2 technical replicates per group; one-way ANOVA). The graph starting point is 1hr post-treatment, when the wound was inflicted. TEER readings at 24h were plotted in a bar graph. Data is represented as % change vs. control (one-way ANOVA, Tukey’s post-test). (**D**) HCMECs were treated with 10uM AβQ22, 1mM Hcy, or a combination of the two for 48h and the samples collected were utilized for an actin polymerization assay (Abcam). Data is represented as % change vs. control (N = 3 experiments with 2 or more technical replicates; one-way ANOVA; Tukey’s post-test). For A-D, **** p<0.0001, *** p<0.001, ** p<0.01, * p<0.05.

Upon binding of VEGF-A to the VEGFR2, the receptor is phosphorylated at tyrosine Y1175 to promote EC proliferation and angiogenesis. Vascular endothelial growth factor A (VEGF-A) and phosphorylated vascular endothelial growth factor receptor 2 (VEGFR2) at tyrosine1175 (pVEGFR2 Y1175) protein expression was assessed by WB in HCMECs treated with Aβ40-E22Q, Hcy, or a combination of both for 6h. Following the combination treatment, we observed an additive decrease in both the expression of VEGF-A (the VEGFR2 ligand), and in pVEGFR2 Y1175 **(Fig. 6A/B)**. To confirm that the combined treatment of HCMECs with Aβ40-E22Q and Hcy inhibits the release of VEGF-A, soluble VEGF-A levels were also analyzed by an ELISA assay in the cell media. Interestingly, HCMECs treated with Hcy or Aβ+Hcy released significantly lower levels of soluble VEGF-A after 24h compared to controls and Aβ-treated cells **(Fig. 6C).**

**Figure 6.**
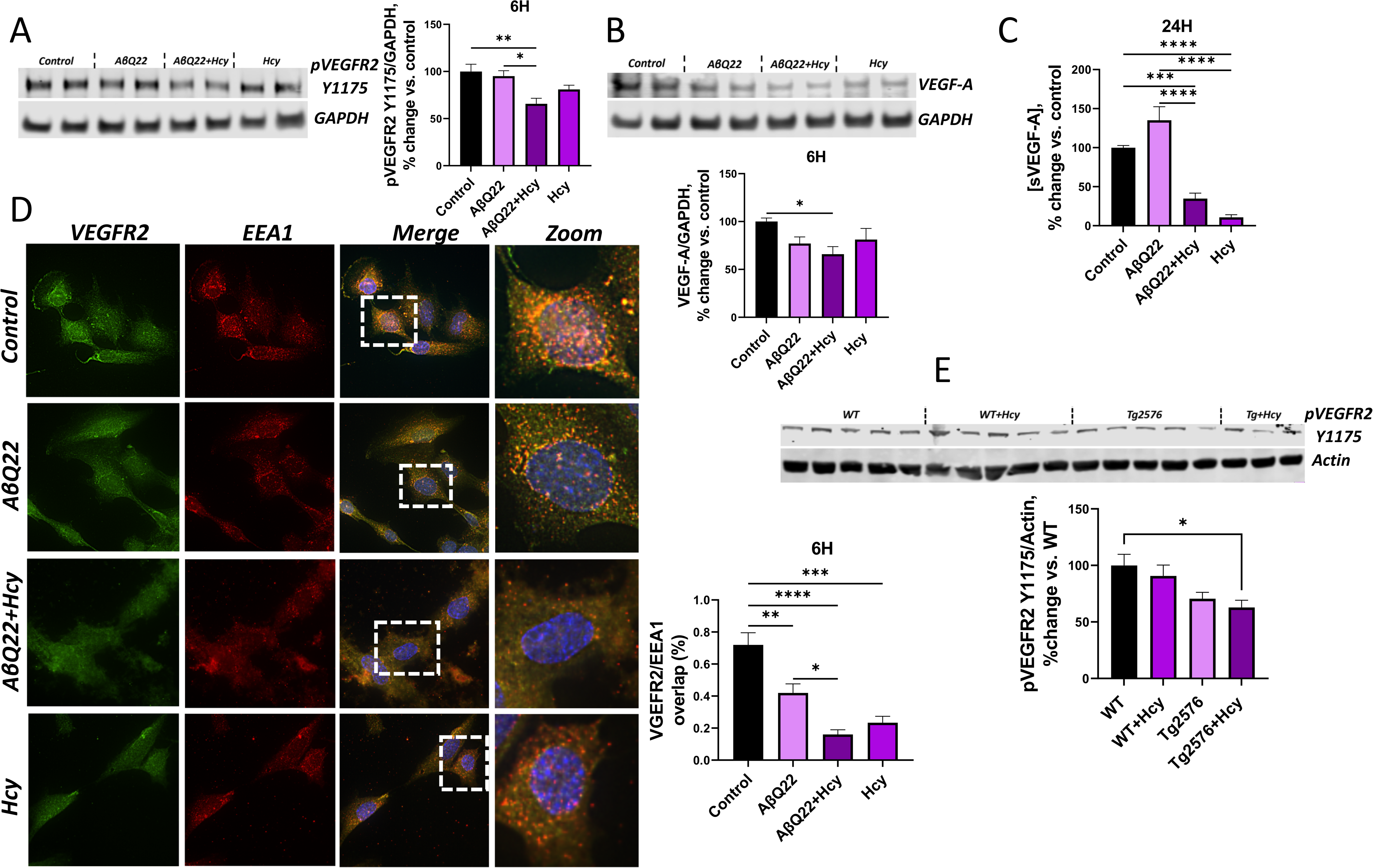
Hcy potentiates Aβ40E22Q-mediated loss of angiogenic mediators and VEGFR2 internalization within early endosomes. (**A**) HCMECs were treated with 25uM AβQ22, 1mM Hcy, or a combination of the two for 6h. pVEGFR2 protein expression was evaluated via western blot and GAPDH was used for normalization. Data is represented as % change vs. control (N = 3 experiments with 2 technical replicates; one-way ANOVA; Tukey’s post-test). (**B**) HCMECs were treated with 25uM AβQ22, 1mM Hcy, or a combination of the two for 6h. VEGF-A protein expression was evaluated via western blot and GAPDH was used for normalization. Data is represented as % change vs. control ((N = 3 experiments with 2 technical replicates; one-way ANOVA; Tukey’s post-test). (C) HCMECs were treated with 25uM AβQ22, 1mM Hcy, or a combination of the two for 24h and media was collected to run a soluble VEGF-A ELISA (sVEGF-A). Data is represented as % change vs. control (N = 3 experiments with 2 or more technical replicates; one-way ANOVA; Tukey’s post-test). (**D**) HCMECs were treated with 25uM AβQ22, 1mM Hcy, or a combination of the two for 6h and VEGFR2 internalization into early endosomes was evaluated by immunocytochemistry. HCEMCs were stained with DAPI (blue), VEGFR2 (green), and EEA1 (red). Colocalization of VEGFR2 fluorescence in EEA1 positive endosomes was analyzed with Halo software (Indica Labs). (**E**) Western blot analysis was utilized to assess pVEGFR2 protein expression in WT and Tg2576 mice -/+ Hhcy. Actin was used for normalization and data is represented as % change vs. control (N= 3 or more mice per group; n=2 technical replicates; one-way ANOVA, Tukey’s post-test). For A-E: **** p<0.0001, *** p<0.001, ** p<0.01, * p<0.05.

Once the VEGFR2 is bound, the receptor dimerizes and is subsequently internalized within endosomes, where it is phosphorylated (Balaji Ragunathrao, Vellingiri, Anwar, Akhter, & Mehta, 2020). Internalization of the phosphorylated VEGFR2 into early endosomes protects the receptor from degradation and sustains the activation of signaling that promotes angiogenesis (Balaji Ragunathrao et al., 2020). To assess how Aβ40-E22Q, Hcy, and the combination modulate VEGFR2 internalization into early endosomes, immunocytochemistry for VEGFR2 and EEA1 (early endosome antigen 1), was conducted in HCMECs treated with 25uM Aβ40-E22Q, 1mM Hcy, or a combination of the two for 6h. In control cells, a prevalent colocalization of VEGFR2 and EEA1 was observed **(Fig. 6D)**. In HCMECs treated with Aβ40-E22Q or Hcy, we detected a notable decrease in both VEGFR2 and EEA1 staining, as well as a decrease in colocalization of VEGFR2 and EEA1. When HCMECs were treated with the combination of Aβ40-E22Q and Hcy, an almost complete loss of VEGFR2 and EEA1 expression was observed, as well as a severe decrease in VEGFR2 and EEA1 colocalization **(Fig. 6D)**. To confirm whether Hhcy contributes to decreased pVEGFR2 Y1175 and potentiates Aβ-mediated pVEGFR2 Y1175 decreases *in vivo*, western blot analysis of pVEGFR2 Y1175 was evaluated in the pre-frontal cortex of WT and Tg2576 mice subjected or not to Hhcy-inducing diet. WT mice with Hhcy and Tg2576 mice demonstrated a trend towards decreased pVEGFR2 Y1175 expression. Importantly, as hypothesized, a significant and additive loss of pVEGFR2 Y1175 protein expression was observed in Tg2576 mice with Hhcy **(Fig. 6E)**.

## Discussion

It is well accepted that cardiovascular risk factors, including Hhcy, increase the risk and pathological severity of AD (Tinelli et al., 2019) (Carey & Fossati, 2023) (Kamat et al., 2015). Previous studies demonstrated that Hhcy accelerates cerebral amyloidosis and CAA, exacerbating deficits in CBF and cerebrovascular dysfunction in AD mouse models (Braun et al., 2019; J. G. Li, Chu, Barrero, Merali, & Pratico, 2014; Zhuo et al., 2010; Zhuo & Pratico, 2010) (J. G. Li & Pratico, 2015) (Sudduth, Powell, Smith, Greenstein, & Wilcock, 2013). Additionally, neuroinflammatory processes, microhemorrhages and cerebrovascular atherosclerosis were observed in post-mortem human brains of patients with Hhcy (Weekman et al., 2022). It is evident that Hhcy has severe pathological consequences on the cerebrovascular environment. However, the specific endothelial molecular mechanisms responsible for these deleterious effects and for the combined impact of Hhcy and amyloid cerebrovascular pathology remained poorly elucidated (Carey & Fossati, 2023). This study demonstrates that a combination challenge of HCMECs with a vasculotropic Aβ peptide, the Dutch mutant, and Hcy exacerbates TRAIL DR4/5-mediated extrinsic apoptosis, BBB permeability, and angiogenesis impairment. Importantly, we described specific concurrent molecular mechanisms through which both Aβ40-E22Q and Hhcy act to induce brain EC dysfunction, and we observed that the combined effect of these two cerebrovascular challenges on cerebral ECs is, in most cases, additive in nature.

Cerebral EC apoptosis contributes to the pathogenesis of AD, sporadic and familial forms of CAA (Take et al., 2022) (X. X. Wang, Zhang, Xia, & Jia, 2020), being one of the main culprits for the appearance of micro and macro-hemorrhages (X. X. Wang et al., 2020). We have previously shown that the vasculotropic Dutch Aβ mutant, Aβ40-E22Q, highly associated with CAA, promotes, TRAIL DRs (DR4/5)-mediated cerebral EC apoptosis (through direct binding to these receptors), BBB permeability, and angiogenesis deficits similarly to Aβ40-WT, albeit with faster kinetics (Fossati et al., 2010; Fossati et al., 2012b; R. Parodi-Rullan et al., 2020; R. M. Parodi-Rullan, Javadov, & Fossati, 2021). Therefore, this peptide represents an excellent tool to study the effects of Aβ-mediated endothelial pathology *in vitro*. Multiple studies have also demonstrated that Hcy promotes cerebral EC dysfunction (Lai & Kan, 2015) (Kamat et al., 2015) (Faraci & Lentz, 2004; Tyagi et al., 2006) (Suhara et al., 2004). However, it is not known if Hcy can induce DR4/5-mediated EC apoptosis.

Our current study is the first to demonstrate that high Hcy potentiates the toxic effects of Aβ on cerebral EC through the DR4/5-mediated apoptotic pathway, and that the combination of Aβ and high Hcy activates this pathway in an additive manner.

We showed that Aβ40-E22Q and Hcy additively increase DR4 and DR5 expression –known to follow these DRs activation (Micheau et al., 1999) (Poh et al., 2007) (Rossin et al., 2009)–, as well as the downregulation of cFLIP, an inhibitor of caspase 8, leading to an additive increase in caspase 8 activation. Activation of caspase 8 leads to both direct caspase 3 activation, as well as Bid cleavage, which results in increased mitochondrial membrane permeability (Kim, Zhao, Barber, Kuharsky, & Yin, 2000), CytC release from the mitochondria, caspase 9 activation, and ultimately caspase 3 activation. Indeed, Aβ40-E22Q and Hcy also produce an additive increase in Bid cleavage, caspase 9 activity and a synergistic increase in caspase 3 activity.

Aβ peptides, including the E22Q variant, are known to promote mitochondria-mediated apoptosis (Fossati et al., 2010; Solesio et al., 2018). Hcy was also found to promote mitochondria-mediated apoptotic pathways within ECs, through a decrease in BCL2/Bax mRNA ratio and an increase in CytC release and caspase 9 activity (Tyagi et al., 2006). Here, we demonstrated that combined treatment of cerebral ECs with Aβ40-E22Q and Hcy results in potentiated decreases in BCL2 and increases in Bax protein expression, which is responsible for the formation of the mitochondrial permeability transition pore (MPTP) (Narita et al., 1998) (Q. Chen et al., 2015) (Kowaltowski, Vercesi, & Fiskum, 2000), increased mitochondrial fragmentation and CytC release. A potentiated CytC release was indeed confirmed by both immunocytochemistry and ELISA, eventually inducing additive increases in DNA fragmentation (apoptosis) and secondary necrosis. Overall, these results demonstrate that all the steps of the DR-mediated apoptotic pathway, which directly triggers mitochondria-mediated apoptosis, are exacerbated in a sequential and additive manner by the combined challenge with Aβ and Hcy. Hence, we revealed that this specific apoptotic pathway may be a potential target for future therapeutic strategies to tackle the cerebrovascular damage induced by both amyloidosis and Hhcy.

Additionally, Aβ peptides, including the Aβ40-E22Q variant, have been shown to decrease TEER, indicative of an increased BBB permeability, (R. Parodi-Rullan et al., 2020). Hhcy can also increase BBB permeability (Beard et al., 2011) (Kamath et al., 2006). This study is the first to demonstrate that combined treatment of HCMECs with Aβ40-E22Q and Hcy potentiates barrier dysfunction, operating in an additive manner on the same molecular mediators of BBB damage. In particular, we showed that Hcy exacerbates the Aβ-mediated decrease in claudin-5 and increased phosphorylation of claudin-5 and VE-cadherin, previously shown to be associated with BBB permeability and retraction of ECs from each other (Yamamoto et al., 2008) (Sidibe & Imhof, 2014) (Hatanaka et al., 2011). Importantly, combined treatment of HCMECs with Aβ40-E22Q and Hcy potentiates an early release of IL2. This cytokine is known to lead to the phosphorylation of VE-cadherin, revealing a specific molecular mechanism through which the combination of Aβ40-E22Q and Hcy potentiates BBB permeability (Wylezinski & Hawiger, 2016). Acute treatment with Aβ40-E22Q and Hcy also additively increased the expression of ICAM, an extravasation molecule that promotes EC recruitment of immune cells into the brain (Dietrich, 2002), as well as MMP2 activation, which results in digestion of the extracellular matrix (Weekman & Wilcock, 2016).

At later time points, we also observed an additive decrease in ZO-1, a vital anchor protein within the BBB, responsible for linking cytoskeletal proteins, such as actin, in one cell, and TJ proteins, such as cadherins, occludins, and claudins, in another adjacent cells (Fanning, Jameson, Jesaitis, & Anderson, 1998; Liu, Wang, Zhang, Wei, & Li, 2012; Tornavaca et al., 2015), as well as for actin filaments organization and polymerization. This finding, which we also confirmed in Tg2576 animals exposed to a Hhcy-inducing diet, explains the dramatic loss of EC barrier integrity and TEER associated with the combined treatment. Accordingly, in patients with comorbid AD/CAA and HHcy, a severe BBB permeability may be expected. This finding is of particular importance in view of the recently FDA approved Aβ-targeted immunotherapies and their probability to induce ARIA (amyloid-related imaging abnormalities), with cerebral edema or hemorrhage, an undesired effect causally mediated by BBB dysfunction. Our data suggests that patients presenting with both amyloidosis (AD/CAA) and Hhcy should be considered at higher risk for these complications, due to their already highly compromised BBB function, and physicians should be cautious in prescribing Aβ immunotherapies in patients with these comorbidities. These findings also highlight the need to continue the search for alternative therapies which could be affective also in patients with cerebrovascular comorbidities (Canepa et al., 2023), which are among the most frequent AD associated comorbidities (Carey & Fossati, 2023).

Angiogenesis is an important repair mechanism, particularly needed when the brain is damaged or hypoperfused, such as in neurodegenerative conditions like AD and CAA. Aβ peptides, including the Aβ40-E22Q variant, have been shown to reduce cerebral EC angiogenic capabilities (R. Parodi-Rullan et al., 2020). Hcy is also known to inhibit angiogenesis (Zhang et al., 2012) (Nagai et al., 2001) in peripheral ECs. This study elucidates Hcy’s effect on cerebral EC angiogenesis and demonstrates that combined treatment of cerebral ECs with Aβ40-E22Q and Hcy potentiates the decreases in angiogenesis progression scores. Furthermore, we revealed that pro-inflammatory cytokines such as IL13, IL4, IL8, IL1β, and IL6, known to induce pro-angiogenic pathways (Ma, Yang, He, Zhang, & Chang, 2021), are additively decreased as early as 3h in EC challenged with both Aβ and Hcy. IL13 and IL4 are both potent inducers of angiogenesis, stimulating the formation of tubular vessels by human microvascular ECs *in vitro*. Specifically, IL13 has been found to promote VEGF-A production (Fukushi, Ono, Morikawa, Iwamoto, & Kuwano, 2000) (Fukushi et al., 1998). IL8 has been shown to regulate angiogenesis by stimulating VEGF expression and autocrine VEGFR2 activation in ECs as well as enhance EC proliferation and survival (Martin, Galisteo, & Gutkind, 2009) (A. Li, Dubey, Varney, Dave, & Singh, 2003). IL-1β potently stimulates angiogenesis *in vitro* within endothelial progenitor cells (Rosell et al., 2009). IL6, a pleiotropic cytokine with pro- and anti-inflammatory functions, has been shown to inhibit angiogenesis in some molecular contexts, but also to stimulate vessel sprouting in ECs and stimulate VEGF production (Gopinathan et al., 2015) (Zegeye, Andersson, Sirsjo, & Ljungberg, 2020). The effect on decreasing these cytokines is therefore a possible upstream mechanism through which Aβ40-E22Q and Hcy additively decrease the activation of angiogenic pathways.

Additionally, this study has been the first to demonstrate that combined challenge with Aβ and Hcy additively diminishes cerebral endothelial barrier wound healing capabilities. More specifically, Aβ and Hcy decrease actin polymerization kinetics, a process important for barrier formation as well as wound healing. Additionally, acute exposure to Aβ and Hcy decreased phosphorylation of VEGFR2 at tryosine1175 (Y1175), that has been associated with EC proliferation and angiogenesis (X. Wang, Bove, Simone, & Ma, 2020), as well as additively reducing the expression of VEGF-A and the release of soluble VEGF-A by HCMECs. Mechanistically, VEGFR2 endocytosis is an essential regulator of angiogenesis signaling. VEGF binding to the VEGFR2 on the EC plasma membrane results in receptor endocytosis into the slow trafficking pathway where the receptor will end up in early endosomes. Internalization within early endosomes promotes VEGFR2 phosphorylation and helps to propagate the downstream signaling and prevent degradation or constitutive recycling of the receptor (Lampugnani, Orsenigo, Gagliani, Tacchetti, & Dejana, 2006). The observed additive decrease in colocalization of the VEGFR2 within early endosomes suggests a significant reduction in VEGFR2 internalization and subsequent pro-angiogenic signaling, and may additionally suggest a potentiated disruption in autophagic mechanisms by Aβ and Hcy, which will need to be further investigated.

Importantly, key mechanisms uncovered by our *in vitro* experiments were confirmed *in vivo* in prefrontal cortex samples of Tg2576 and WT mice subjected to Hhcy-inducing diet. WT mice with Hhcy and Tg2576 mice demonstrated trends towards increased DR4 expression and decreased ZO-1 and pVEGFR2 Y1175 expression. However, in Tg2576 mice with Hhcy, an additive and significant upregulation of DR4 expression, downregulation of ZO-1 and reduction in pVEGFR2 Y1175 were observed. These results provide *in vivo* evidence of Hhcy’s ability to potentiate Aβ-induced DR-mediated apoptosis, BBB dysfunction, and angiogenic downregulation by modulating the same molecular mechanisms revealed by our HCMECs results.

This study is not free of limitations. Mimicking the chronic condition of Hhcy *in vitro* presents challenges. Individuals with Hhcy have Hcy level from 15uM to over 100uM concentrations, circulating throughout their blood and organs for years. To mimic acutely, for short treatments *in vitro,* the effects of the chronic physiological exposure to Hhcy, most previous studies on ECs utilized concentrations of 1mM or higher, necessarily higher than physiological levels, with 1mM being the initial concentration that activated endothelial apoptosis in most reports (Nagai et al., 2001) (Fan et al., 2019) (J. Chen, Huang, Hu, Bian, & Nian, 2021) (Jia, Lai, Zhao, Gong, & Zhang, 2013). Considering that our *in vitro* treatments span from a few hours-2 days, we decided to also utilize for most experiments a 1mM concentration, the lowest concentration shown to induce apoptosis in previous EC studies, for our HCMECs. However, the corroborating results of our mouse brain experiments, where chronic treatment with Hhcy is performed for months at non-lethal concentrations, serve to confirm the physiological validity of our *in vitro* data.

Additionally, future studies will be needed to investigate Hhcy’s ability to potentiate Aβ-induced DR-mediated apoptosis, BBB dysfunction, and angiogenesis deficits in AD/CAA animal models and in human AD/CAA brains longitudinally with disease progression, in different cerebral regions and within isolated cerebral vessels. Finally, other pathways, such as autophagy or ER stress, may also be involved in the combined effects of Aβ and Hcy, and future research endeavors will be necessary to tackle these additional mechanisms.

## Conclusions

Overall, this study demonstrates that the combined exposure of cerebral EC to Hcy, a potent CV risk factor for dementia, and to a vasculotropic Aβ peptide additively promotes TRAIL-DR mediated apoptosis, induces BBB dysfunction and decreases cerebral EC repair mechanisms and angiogenic impairment, by acting on the same molecular mechanisms within each pathway. EC death and barrier dysfunction lead to a leaky BBB, resulting in microhemorrhages, infiltration of peripheral immune cells, neuroinflammation as well as decreased cerebral microvessel density and hypoperfusion. Angiogenesis and cerebrovascular wound healing, vital compensatory mechanisms capable of regenerating and repairing the brain microvasculature, are also additively damaged by the two challenges. Hence, this work suggests that the comorbid presence of Hhcy in AD and CAA patients will potentiate the vicious cycle of Aβ-induced endothelial and BBB damage and inability to repair this damage, resulting in hypoperfusion, neurovascular unit dysfunction and the resulting poor clearance of amyloid, therefore contributing to a vicious cycle of increased amyloid deposition and toxicity. Ultimately, this study identified specific endothelial pathways that are activated or disrupted in an additive manner by Hcy and Aβ to produce AD-related cerebral microvascular pathology, revealing novel molecular targets for AD and CAA therapy. This work also paves the way to establishing possible preventive or therapeutic strategies to reduce the risk of dementia in individuals with CV risk factors such as Hhcy.

## List of Abbreviations

AD: Alzheimer’s Disease
EC: Endothelial Cell
BBB: Blood Brain Barrier
CV: Cardiovascular
Hhcy: Hyperhomocysteinemia
Hcy: Homocysteine
Aβ40: Amyloid-beta 40
TRAIL: TNF-related apoptosis-inducing ligand
DR4/DR5: Death Receptor 4/5
CAA: Cerebral Amyloid Angiopathy
CBF: Cerebral Blood Flow
TJ: Tight Junction
ZO-1: Zona Occludin 1
CytC: Cytochrome C
HCMECs/D3: Human Cerebral Microvascular Endothelial Cells/D3
HCECs: Primary Human Cerebral Endothelial Cells
WT: Wild-type
TEER: Trans-endothelial electrical resistance
VEGF-A: Vascular Endothelial Growth Factor A
VEGFR2: Vascular Endothelial Growth Factor Receptor 2
EEA1: Early Endosomal Antigen 1
MPTP: Mitochondrial Permeability Transition Pore

## Declarations

### Ethics approval and consent to participate

Not Applicable

### Consent for publication

Not Applicable

### Availability of Data and Materials

The data included in this article will be shared upon reasonable request to the corresponding author and will be made available after publication in data biorepositories.

## Funding/Acknowledgements

This work was supported by NIH R01NS104127 and R01AG062572 grants, the Edward N. and Della L. Thome Memorial Foundation Awards Program in Alzheimer’s Disease Drug Discovery Research, the Alzheimer’s Association (AARG-20-685663), the Pennsylvania Department of Heath Collaborative Research on Alzheimer’s Disease (PA Cure) Grant, awarded to SF, and by the Karen Toffler Charitable Trust, and the Lemole Center for Integrated Lymphatics research.

## Authors Contributions

AC and SF designed the study. AC conducted most of the experiments in this manuscript and wrote the manuscript. SF provided oversight and scientific insight on experimental planning, assisted and mentored AC in writing, edited the manuscript and obtained funding for this project. RPR conducted the ECIS TEER experiment. RVT bred and maintained the mouse cohorts utilized in this study and assisted with imaging and quantifying immunocytochemistry experiments. EC assisted with running western blots with mice samples. All authors read and approved the final manuscript.

---

**Supplementary Figure 1.**
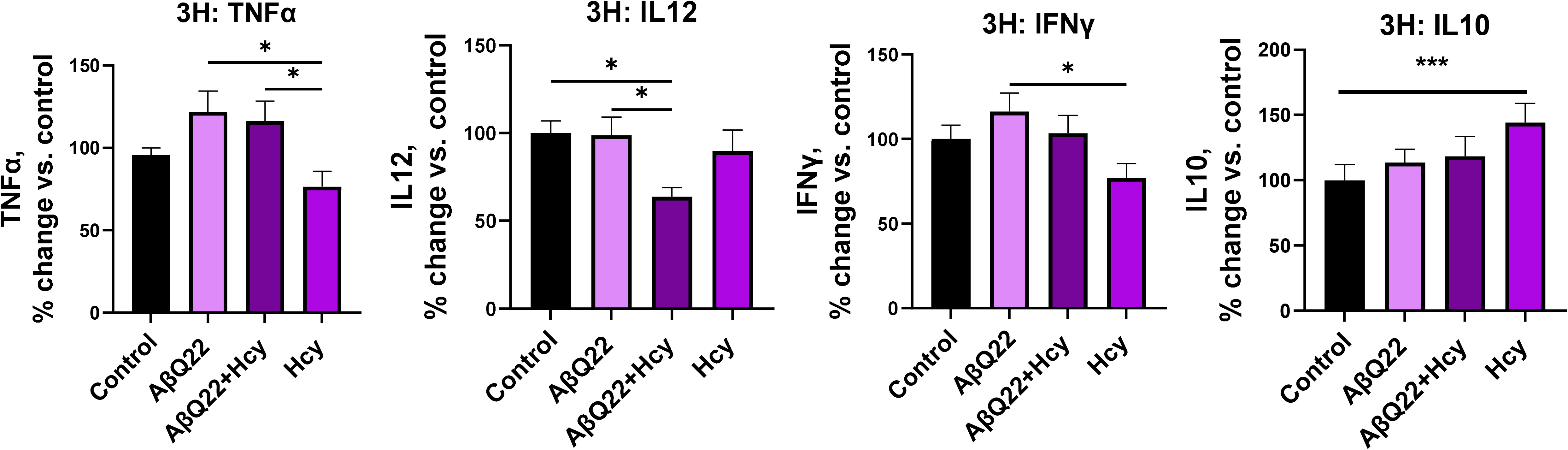
Effects of Hcy and AβQ22 treatment on other pro-inflammatory cytokine release in HCMECs media. HCMECs were treated with 25uM AβQ22, 1mM Hcy, or a combination of the two for 3h and media was collected to run a multiplex inflammatory cytokine assay (MesoScale Discovery). Protein concentration was used for normalization. Data is represented as % change vs. control (N ≥ 3 experiments with 2 or more technical replicates; one-way ANOVA; Tukey’s post-test).Hcy significantly reduced TNFα and increased IL10 release compared to control cells. The combination of AβQ22 + Hcy decreased the release of IL12 compared to Cnt and AβQ22-treated cells.

